# Studying of RNA-protein interactions between a mutant form of glycyl-tRNA synthetase associated with neurodegenerative diseases and IRES type I

**DOI:** 10.1101/2023.03.29.534367

**Authors:** E. S. Vinogradova, A. O. Mikhailina, O. S. Nikonov, E. Yu. Nikonova

## Abstract

Aminoacyl-tRNA synthetases (aaRS) are the main enzymes of protein biosynthesis. Human glycyl-tRNA synthetase, in addition to the main function of amino acid transfer to the corresponding tRNA molecules, is also involved in the initiation of IRES I translation. All members of the enterovirus genus have this type of IRES.

It is also known that the presence of point mutations in aaRS leads to the occurrence of diseases in which peripheral nerves are affected. One such disorder of the nervous system is the incurable neurodegenerative disorder Charcot-Marie-Tooth (CMT). The most studied enzyme whose mutations cause CMT is glycyl-tRNA synthetase (GlyRS).

In this work, we tested the ability of various mutant forms of glycyl-tRNA synthetase associated with Charcot-Marie-Tooth syndrome to form a stable complex with IRES I. It turned out that neither catalytic activity nor the ability to form a dimer are necessary for the interaction of GlyRS with IRES.

## Introduction

Human glycyl–tRNA synthetase (GlyRS) consists of three domains: a WHEP domain, a catalytic domain, and an anticodon-binding domain.

The WHEP domain (short for the names of synthetases containing this domain: TrpRS –W, HisRS –H and GluProRS –EP) is located at the N–terminus of the protein, has a helix–turn–helix conformation, is a non–specific RNA-binding domain, interacts with DNA and participates in protein–protein interaction (Qin *et al*., 2014). Removal of the WHEP domain does not affect the enzymatic activity of tRNA synthetases, indicating that the role of this domain is not related to aminoacylation. The catalytic domain is formed by eight β–strands surrounded by α-helices on both sides. (Deng *et al*., 2016). This domain also insist of 3 motifs, where motif 1 participates in dimer formation; motifs 2 and 3 include conserved amino acid residues with charged and polar side chains involved in the recognition of substrates: glycine and ATP (Qin *et al*., 2014).

The anticodon–binding domain is located at the C–terminus of the glycyl–tRNA synthetase. It is a globular α/β domain typical for IIa class synthetases. It is responsible for the specificity of binding to the nucleotides of the anti-codon tRNA.

In 2012, a new non–canonical function of human glycyl–tRNA synthetase was discovered - regulation of translation initiation in polioviruses. It has been shown that human GlyRS is able to bind to the V domain of the poliovirus IRES (whose apical part mimics the anti-codon hairpin tRNAGly) and enhance translation initiation (Andreev *et al*., 2012). However, the exact mechanism of this process has not yet been studied in detail.

A connection of such a severe neurodegenerative disease as Charcot–Marie–Toute disease with missense mutations in the glycyl–tRNA synthetase gene was reported for the first time in 2003. (Antonelli et al, 2003). This disease is the most common hereditary motor–sensory neuropathy in humans. Since this disease is associated with point mutations in *GARS*, pathological variants of glycyl–tRNA synthetase are fairly well characterized biochemically.

To date, more than 20 mutations associated with various neurodegenerative diseases have been described in the *GARS*. Substitutions L129P and G240R are the most common mutation variant. For us, these substitutions are interesting because they are located in the inter-domain interface of GlyRS (Fig. 1). It is known from the literature that these mutations disrupt dimer formation, and such mutant proteins are completely catalytically inactive. Since the dimeric form of GlyRS is necessary for aminoacylation, the loss of the catalytic activity of the enzyme with L129P and G240R substitutions is clearly associated with the destruction of the dimer (Nangle et. al., 2007).

**Figure 1.**
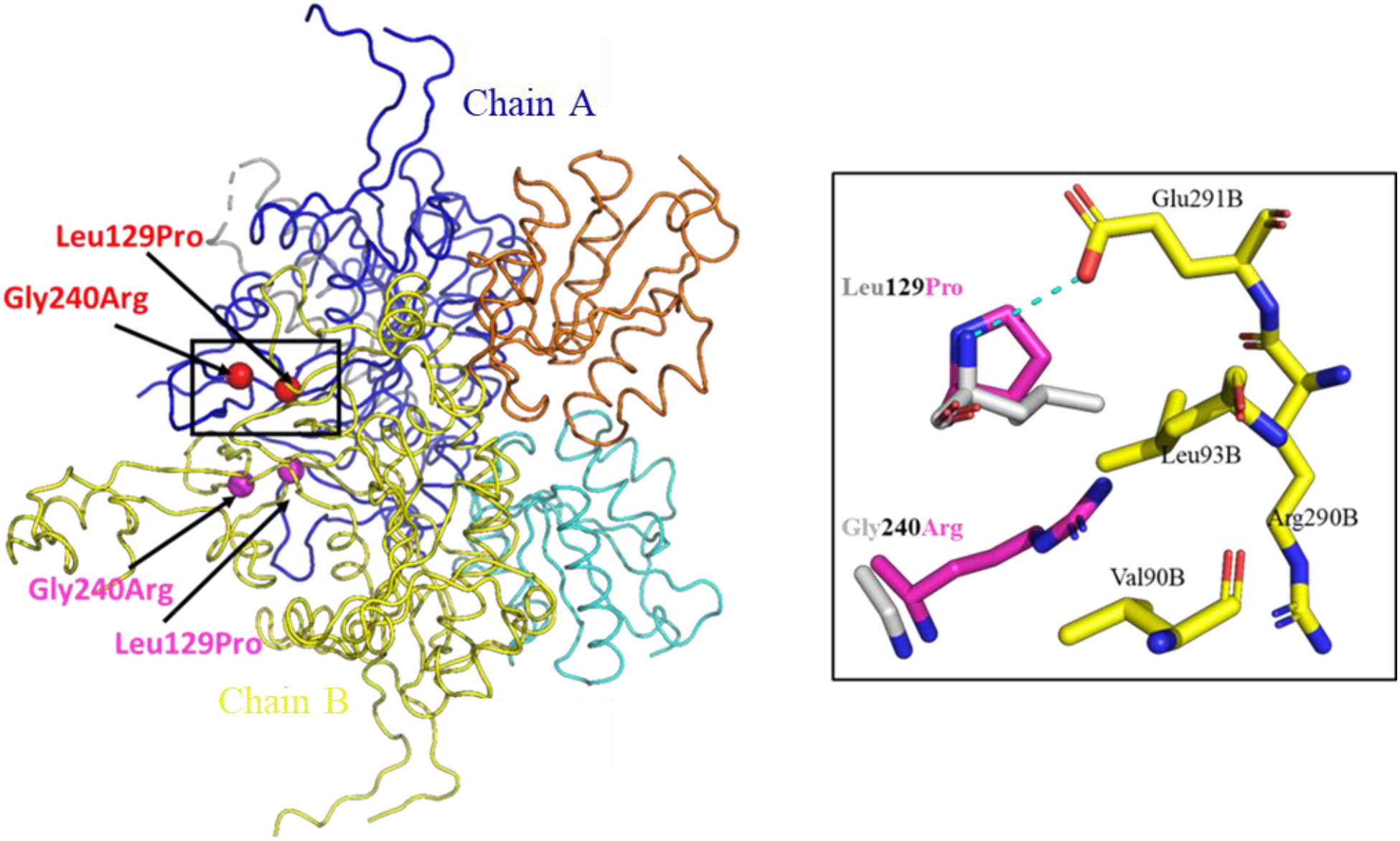
Spatial structure of human glycyl–tRNA synthetase (PDB ID: 5E6M). Chain A: gray indicates the WHEP domain; blue indicates the catalytic domain; turquoise indicates the ABD domain; red balls show the position of 129 and 240 amino acid residues; Chain B: yellow indicates the catalytic domain; orange indicates the ABD domain; fuchsia balls indicate the position of L129P and G240R in chain B.

For our work, these mutant forms of glycyl-tRNA synthetase were sutable because they allowed us to check whether a dimeric form of human glycyl–tRNA synthetase is necessary for the formation of a stable complex with IRES type I and whether the GlyRS monomer was capable to bind IRES.

The position of GlyRS missense mutations affecting dimer formation are shown in the black square: gray – GlyRS WT; fuchsia is a mutant form of glycyl–tRNA synthetase: yellow is the position of the amino acids of the chain A

## Methods

### Introduction of mutations into the gene of human glycyl-tRNA synthetase; isolation of proteins in preparative quantities

The previously obtained pET-22b–GARS ΔWHEP plasmid (Nikonova *et al*., 2016) was used as a matrix for mutations introduction (GlyRS L129P; GlyRS G240R and GlyRS L129P/G240R).

Mutations were introduced using the Quick change method (Liu *et al*., 2008)

Primers used for PCR:

GlyRS L129P forward: 5’–CGATTGCACCATGCCGACCCCTGAGCCAG–3’
GlyRS L129P revers 5’–CTGGCTCAGGGGTCGGCATGGTGCAATCG–3’
GlyRS G240R forward 5’–CTGGAGGAAACATGCCTAGATACTTGAGACCAGAAACTG–3’
GlyRS G240R 5’–CAGTTTCTGGTCTCAAGTATCTAGGCATGTTTCCTCCAG–3’

After all the substitutions were made, the absence of undesirable mutations was checked by sequencing.

The genes of the mutant forms of glycyl-tRNA synthetase were expressed in *E. Coli* BL21(DE3) cotransformed by the Rosetta plasmid (Novy *et al*., 2001). The cell culture was grown at 37 °C with intensive stirring (180 rpm) on a rich TB medium with the addition of selective antibiotics: ampicillin and chloramphenicol, up to an optical density of D590 = 0.8. Isopropyl-β-D-1-thiogalactopyranoside (IPTG) was added to a final concentration of 0.5 mM. After the addition of the inductor, the cells were incubated at 20 ° C for 17 hours.

### Isolation of recombinant proteins

The cells of the superproducer strains were resuspended in a buffer containing 50 mm Tris–HCl, pH 9.0, 500 mM NaCl, 5mM β–ME, 1 mM PMSF, 0.1% Triton X-100 and disrupted using a high–pressure homogenizer Avestin (Canada) The lysate was centrifuged for 30 min at 14,000 g and 4°C. The supernatant was applied to a column with HisTrap ™ High Performance resin (5 ml) for affine chromatographic purification. After applying the samples, the column was washed with 6 column volumes with a buffer of 50 mM Tris–HCl, pH 9.0, 2 M LiCL, 5mM β–ME, 0.1% Triton X–100. Then 6 column volumes with a buffer of 50 mM Tris–HCl, pH 9.0, 50mM NaCl, 5mM β–ME, 0.1% Triton X–100. Elution was performed with a linear gradient (20-450 mM) of imidazole in a buffer of 50 mM Tris–HCl, pH 9.0, 50mM NaCl, 5mM β–ME, 0.1% Triton X–100. The volume of the linear gradient is 120 ml, the volume of the fraction is 4 ml.

Fractions containing target proteins were combined and applied to a column with Q–Sepharose Fast Flow’ (5 ml) balanced with a buffer of 50 mM Tris–HCl, pH 9.0, 50 mM NaCl, 5mM β–ME, 0.1% Triton X–100. For protein elution, a sodium chloride gradient from the starting buffer to 550 mM was used. For the final purification of the preparations, gel filtration was used on HiLoad 16/60 Superdex 200 resin, balanced with a buffer of 50 mM Tris–HCl, pH 9.0, 150 M NaCl, 5mM β–ME, 0.1% Triton X–100. Fractions containing the purified protein were concentrated using a centrifuge concentrator with a molecular weight cutoff of 50 kDa. The purity of the sample was evaluated electrophoretically in 15% PAAG in the presence of SDS.

### Obtaining of IRES fragments

IRES fragments were obtained as described earlier (Nikonova *et al*., 2018).

### RNA-protein complexes formation

Formation of RNA-protein complexes was carried out the same way as described earlier for wild type of glycyl–tRNA synthetase (Nikonova *et al*., 2016).

### Kinetic experiments

Kinetic experiments were conducted as described earlier (Nikonova *et al*., 2016).

## Results and discussion

The level of glycyl-tRNA synthetase production using LB medium was quite low. The increase in the expression level of the target genes was achieved by changing the microbiological medium from LB to TB (Terrific Broth). TB supports the continuous growth of *E. coli* cells in the log phase, providing a high cell density. As a result, the yield of recombinant proteins is significantly increased. The changing the medium led to a 4-fold increase in the yield of pure protein (1.6 mg of pure protein from 1 gram of cell culture grown on LB, and 6.4 mg of pure protein from 1 gram of cell culture grown on TB).

The cytoplasmic form of glycyl-tRNA synthetase consists of three domains: an unstructured WHEP domain, a catalytic core domain, and an anti-codon-binding ABD domain. It was previously shown that the ability to stimulate translation in both a full-sized enzyme and its variant with a cut-off WEB domain is the same (Andreev *et al*., 2012). This was also confirmed by kinetic experiments: GlyRS△WHEP interacts with a fragment of the poliovirus IRES element in the same way as a full-size glycyl-tRNA synthetase, which gives us the opportunity to use a shortened version of the protein for our experiments (Nikonova *et al*., 2016).

Mutant forms (GlyRS Δ WHEP G240R and GlyRS Δ WHEP L129P/G240R) are strongly prone to aggregation. Both proteins are in solution in the form of multimeters.

It was detected by gell-filtration and DLS (data not shown).

To estimate affinity of glycyl-tRNA synthetase and its mutant forms to the fragment of IRES I, we conducted kinetic experiments using the surface plasmon resonance method. Determined kinetic constants are shown in Table 1.

**Table 1.** Dissociation constants of complexes of mutant forms of human glycyl-tRNA synthetase with the fragment of IRES I

| Complex | KD, nM |
| --- | --- |
| GlyRS Δ WHEP + CDV2 | 9,27±2,5 |
| GlyRS Δ WHEP L129P + CDV2 | 4,85±1,1 |
| GlyRS Δ WHEP G240R + CDV2 | - |
| GlyRS Δ WHEP L129P/G240R + CDV2 | - |

This experiment showed that the mutant form of glycyl–tRNA synthetase GlyRS Δ WHEP L129P binds the IRES fragment only two times worse than the control protein GlyRS Δ WHEP, which is not a significant difference. I.e., the catalytically inactive monomeric form is able to form a specific stable complex with IRES, as well as the dimeric form of the protein. Therefore, it can be concluded that the ability to form a dimer is not a necessary condition for the GlyRS–IRES interaction. For mutant forms of GlyRS Δ WHEP G240R and GlyRS Δ WHEP L129P/G240R, we were not able to measure binding constants. This is most likely due to the fact that these mutant forms are strongly prone to aggregation. The main contribution to the aggregation process is made by a mutation in the 240 position. When the monomers of these mutant forms stick together, it seems that the RNA binding site is masked and, as a result, the interaction disappears (is not detected).

Most likely, exact this tendency to aggregation of these mutant forms of glycyl-tRNA synthetase leads to the occurrence of neurodegenerative disease.

## Acknowledgments

We express our gratitude to A. Kazakov (IBP RAS, Pushchino, Moscow region) for his assistance in conducting kinetic experiments on the basis of the Center for Collective Use of IPB RAS.

The work was carried out with the financial support of the Russian Foundation for Basic Research (grant No. 19-34-90135).

## Notes

### Competing Interest Statement

The authors have declared no competing interest.

